# Detection of whole-field luminance changes by superficial interneurons in the zebrafish tectum

**DOI:** 10.1101/178970

**Authors:** Alison J. Barker, Thomas O. Helmbrecht, Aurélien A. Grob, Herwig Baier

## Abstract

The superficial interneurons, SINs, of the zebrafish tectum, also known as tectal horizontal cells, have been implicated in a range of behavioral responses to visual cues, including prey capture and looming-evoked escape. However, there is limited information on their tuning properties, synaptic connectivity and molecular diversity. We have begun to close this gap in our knowledge by performing functional GCaMP6 imaging and electrophysiological recordings of SIN responses to simplified stimuli. We found that a large subset of the SINs strongly responded to luminance changes, with a preference for OFF steps. Although SINs were generally tuned to any dark-light, or light-dark, transition, they were often most sensitive to a narrow range of luminance steps. Tuning properties did not vary with the position of the SIN cell body along the depth of the tectal neuropil or with the orientation of the cell’s neurites within the neuropil. Receptive field (RF) mapping revealed that SINs sample over large areas of the visual field (>20°, typically 50°). RFs of individual SINs were sometimes spatially incoherent, indicating that SINs may receive sparse inputs from multiple RGCs scattered widely over the retina. Lastly, we found that, while most SINs express GABAergic markers, as reported before, a substantial minority express a vesicular glutamate transporter. We propose that the role of the SINs is more complex than previously reported, and provide evidence that at least one SIN subtype could function as a multi-purpose circuit element for signaling changes in ambient luminance for subsequent processing in the tectum.

## Introduction

Ensuring behavioral reliability across variable environmental conditions requires our brains to extract relevant features from the visual landscape, a process which is refined over multiple levels of visual processing. Yet, the exact synaptic and cellular-level mechanisms underlying these transformations remain unknown. Recently the larval zebrafish has emerged as a highly tractable model for studying visual behaviors and interrogating underlying neural circuits (Barker and Baier, 2015; Dunn et al., 2016; Kubo et al., 2014; Dal Maschio et al., 2017; Orger et al., 2008; Portugues and Engert, 2009; Portugues et al., 2014; Semmelhack et al., 2014; Temizer et al., 2015; Thiele et al., 2014).

In the zebrafish brain, the optic tectum is the main center for visual processing. Efforts to map the circuitry of the optic tectum have been limited by methods for reliable identification of its cellular components (Nevin et al., 2010). Despite these limitations, the organization of visual information arriving in the tectum is relatively well-defined: 1) Nearly all retinal ganglion cells (RGCs) send axonal projections into the tectum, either directly or by first sending collaterals to extratectal arborization fields. 2) Retinal axons within the tectum are highly structured, displaying both retinotopic and laminar organization, which are fixed early in development (Robles et al., 2013, 2014). In contrast, the organization of resident cell types within the tectum has only been determined in a very broad sense. The majority of cell bodies reside in the stratum periventriculare (SPV), a dense collection of periventricular neurons (PVNs) below the neuropil. Subclasses of PVNs project outside of the tectum (PVPNs) while interneurons (PVINs) restrict both their axonal and dendritic arbors to the tectal neuropil (Nevin et al., 2010; Robles et al., 2011). A third, much less abundant type of tectal neuron, named the superficial interneuron (SIN) is unique in that its cell body is positioned in the most superficial layers of the tectal neuropil (Del Bene et al., 2010; Nevin et al., 2010).

Due to their spatial segregation, the SINs have provided a unique foothold for dissecting visual processing in the optic tectum. The sparseness of cell bodies in the neuropil has allowed for targeted electrophysiology, functional imaging and ablation. Somewhat surprisingly, the SINs have been implicated in a wide range of behaviors, from prey capture to avoidance (Barker and Baier, 2015; Del Bene et al., 2010; Dunn et al., 2016; Preuss et al., 2014). The breadth of visual behaviors associated with the SINs suggests a global role in visual processing or the existence of functional subsets with specific contributions to diverse visually mediated behaviors. A more comprehensive classification of the SINs is needed to resolve this matter. Here we undertake this task using molecular, functional imaging and electrophysiological techniques. We find evidence of a global role for SINs in the detection of ambient luminance changes, and also discover unanticipated heterogeneity in marker expression, morphology and transmitter use of this important cell class.

## Material and Methods

### Zebrafish care and maintenance

Zebrafish (*Danio rerio*) were maintained at 28°C on a 14hr light/10hr dark cycle following standard procedures (Westerfield et al., 2009). All animal procedures were performed in accordance with regulations set forth by the Max Planck Society and the Regierung von Oberbayern. The following transgenic lines were used and maintained in a TLN background (*Tüpel long-fin* (TL) wild-type strain carrying mutations in *mitfa* (nacre, N)): *Et(Gal4-VP16)s1156t, Tg(UAS:Kaede)*^*s*^1999^*t*^*, Tg(UAS:RFP, cry:EGFP), Tg(UAS:GCaMP6s)mpn101, Tg(Isl2b.2:Gal4-VP16)*^*zc*^65^^, *Tg(14xUAS:EGFP),* and *Tg(UAS:mCherry)s1984t*. The *Tg*(*LCR*^*RH2*^*-RH2-2:GFP*)^*pt115-c*^ line was obtained from X. Wei (Fang et al., 2013). *Tg(vglut2a:loxP-DsRed-loxP-GFP)* and *Tg*(*gad1b*: *loxP-DsRed-loxP-GFP*) lines were acquired from S. Higashijima (Satou et al., 2013). The *reelin:Gal4, UAS:RFP* line was acquired from F. Del Bene (Auer et al., 2014). All fish used in experiments were 6-10 days post fertilization (dpf).

### Enucleation experiments

Bilateral eye removal was performed one to two days prior to imaging experiments. Larvae were anesthetized with 0.02% tricaine and embedded in 2% low melting point agarose (Invitrogen). Eyes were removed surgically with a sterile 25G needle and #55 forceps (Fine Science Tools). Following the procedure larvae were removed from agarose and allowed to recover overnight. The overall health of the larvae was assessed before continuing to imaging experiments.

### Immunostaining

Larvae (8dpf, *Gal4s1156t, UAS:EGFP*) were fixed in 4% PFA in PBS overnight at 4°C followed by a 12 hour incubation in 30% sucrose in PBS. Larvae were embedded in O.C.T. (Tissue-Tek) and sectioned into 12 µm sections. Sections were allowed to dry overnight and subjected to the following immunostaining protocol: 2 minute rehydration in PBS, 30 minute incubation in 0.1% sodium citrate buffer, a brief wash in PBS (∼2 minutes), 5 minute RT incubation in 1% SDS, three 15 minute washes in PBS, a one hour RT incubation in blocking solution (3% Donkey Serum (Sigma-Aldrich) and 0.3% Triton X-100 in PBS) and an overnight primary incubation (Mouse anti-Reelin (1:500 dilution, Calbiochem #553730) and Chicken anti-GFP (1:1000 dilution, Invitrogen A10263)) in blocking solution at 4°C. The following day sections were washed three times with PBS (∼15 minutes per wash) and incubated in secondary antibody (1:200, Invitrogen, Goat anti-Mouse IgG (H+L), Alexa Fluor 546 (A-11030) and Goat anti-Chicken IgY (H+L), Alexa Fluor 488 (A-11039) diluted in blocking solution. Sections were washed three times with PBS (∼15 minutes per wash) and mounted in Fluoromount (Sigma-Aldrich) prior to imaging.

### Statistics

All statistical tests were performed using GraphPad Prism 6.0. When appropriate unpaired t-tests or one and two-way ANOVA tests with Tukey’s correction for multiple comparisons were performed. In all figures * denotes p < 0.05, ** denotes p < 0.005, and *** denotes p < 0.0005, n.s. denotes not statistically significant. All error bars are s.e.m.

### Confocal imaging

Confocal images were acquired with Zeiss LSM700 or LSM780 microscopes with associated ZEN software. For all figure images a minimum of 4 larvae were imaged per condition.

### Colocalization analysis

Colocalization analysis was performed using the Coloc2 custom plug-in for ImageJ (Schneider et al., 2012). In brief, corresponding z stacks for green and red fluorescence were split. ROIs for each GFP + cell (SINs) were defined and colocalization analysis run for both channels within the ROI in the sub-stack where the GFP+ cell body was present. A Pearson’s R value coefficient for colocalization of pixel intensity within each ROI was generated. Pearson coefficients of 0.2 and greater were counted as colocalized for colocalization analysis in crossed transgenic lines and 0.5 and greater for colocalization of immunostainings.

### Calcium imaging and analysis

Calcium imaging was performed using either a custom-built moveable objective 2-photon microscope (MOM, Sutter Instruments) with ScanImage software for image acquisition (Pologruto et al., 2003) or a commercial 2P microscope (Femtonics 3DRC, Femtonics, Hungary and see (Dal Maschio et al., 2017)) with associated software for image acquisition. Both microscopes used 20x objectives (Olympus) and 920 nm excitation for GCaMP6s. Prior to imaging experiments larvae were embedded in 2.5% low melting point agarose (Invitrogen). All larvae were raised on a normal light/dark cycle and kept in a dark room prior to and during imaging (∼0.2 lux). Imaging acquisition was performed between 2.94 -3.37 Hz. For analysis of whole-field flash and looming stimuli calcium imaging data, raw fluorescence traces were analyzed using the Time Series Analyzer V3.0 plug-in for Fiji (Pologruto et al., 2003) as previously described (Barker and Baier, 2015). In brief, ROIs were manually defined for all cell bodies and ΔF/F (ΔF/F = (F_t_ - F_0_)/F_0_, where F_t_ is the fluorescence at time t and F_0_, the baseline fluorescence) values calculated after background subtraction of stimulus presentation artifacts. Cells were excluded from analysis if they did not show responses across two trials, did not have a minimum ΔF/F response of 0.8 for all trials or if the baseline (40 frames) before the onset of the visual stimuli had a ΔF/F value greater than 0.2. Responses to stimuli were taken as an average of the four frames immediately following stimulus offset. Responses were averaged across trials for each fish and then across individual fish. 57 cells (47 SINs) in 17 *Gal4s1156t, UAS:GCaMP6s* larvae were analyzed in Figure 1, and 2. Six of these cells were also tested with looming stimuli.

**Figure 1:**
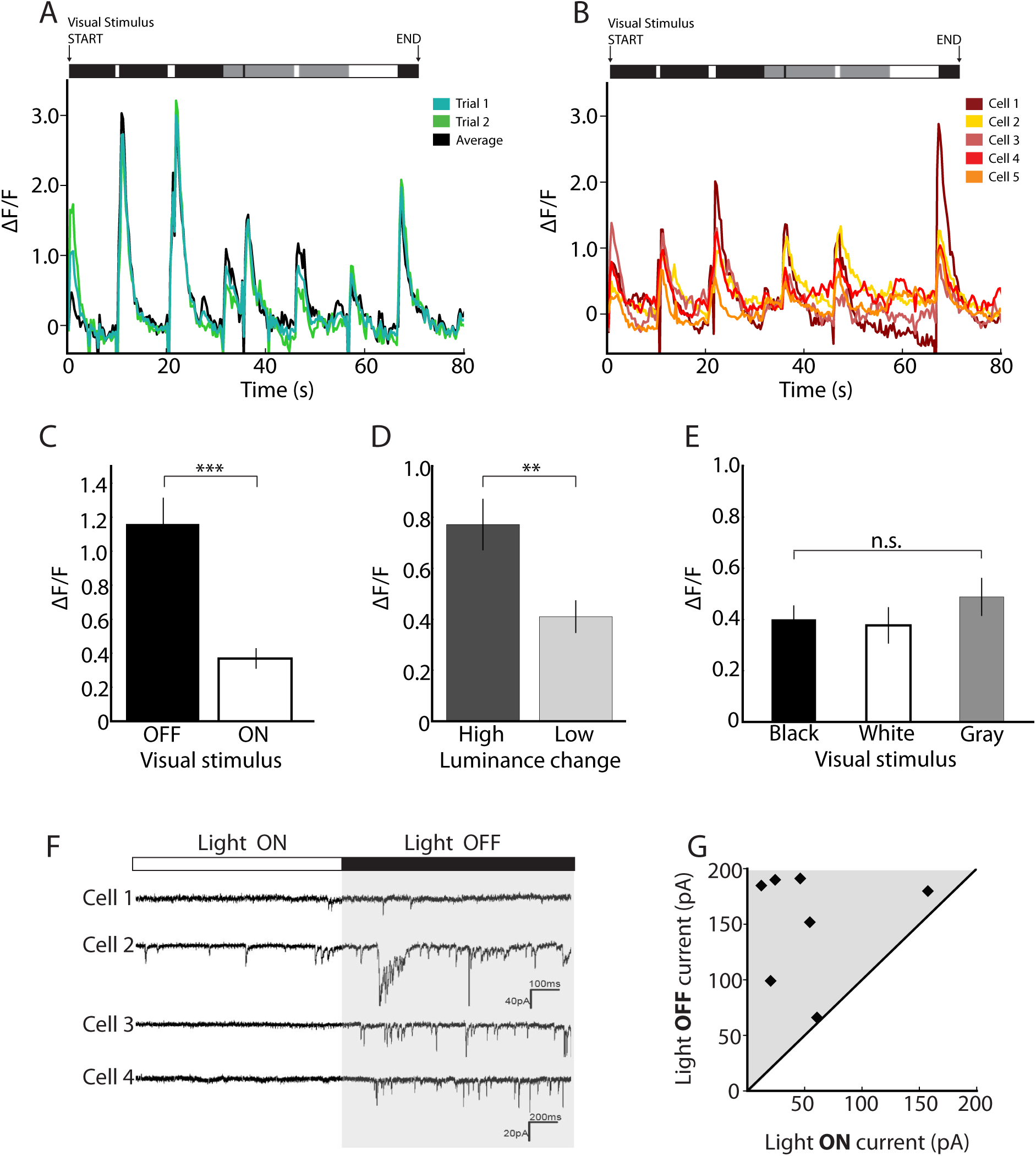
SINs are strongly tuned to changes in whole-field luminance. (**A**) Example cell (7 dpf) in a *Gal4s1156t*, *UAS*:*GCaMP6s* larva shows consistent responses to all whole-field changes in luminance. Visual stimulus is shown above the trace. (**B**) SIN responses to whole-field luminance are highly consistent across larvae. Here 5 cells from four *Gal4s1156t*, *UAS*:*GCaMP6s* larvae show robust responses to whole-field luminance stimuli. (**C**) SINs are more responsive to darkening (OFF) whole-field flashes than brightening (ON) whole-field flashes (unpaired t-test, p < 0.0001) (**D**) SIN responses are greater when the luminance change is high (black to white or white to black) when compared to a low luminance change (white and black transitions to and from gray) (unpaired t-test, p = 0.003). (**E**) SIN responses are specific to transitions between overall luminance levels. No differences in calcium responses are observed during sustained presentation of whole-field black, gray and white stimuli (one-way ANOVA, Tukey’s correction, p = 0.468). (**F, G**) Electrophysiological recordings show increased excitatory synaptic input in response to transitions between light ON and light OFF conditions. (**F**) Four example traces from neurons targeted in the *Gal4s1156t*, *UAS*:*Kaede* line. EPSCs are more frequent under recording conditions of low luminance (Light OFF). (**G**) Total excitatory postsynaptic current is plotted for OFF and ON light conditions. Cells show greater responses under light OFF conditions. For C-E n = 47 cells, 17 larvae. *** denotes p < 0.0005; ** p< 0.005, * p < 0.05, n.s. = not statistically significant. Scale bar in (F), 20 pA, 200 ms. Error bars are s.e.m.

### Visual stimulus presentation

Visual stimuli were presented using an OLED screen (eMagin) with three magenta filters (Figure 1, 2, 6) or a portable LED projector using only the red channel (LG, Model No. PA72G), Figure 3, 4). Presentation of looming stimuli was performed using an expanding disk of constant radial velocity (expanding at 20°/s) as in Temizer et al., 2015. Stimuli for full-screen flash and looming experiments were generated with a custom programmed graphical interface for Vision Egg (Straw, 2008).

**Figure 2:**
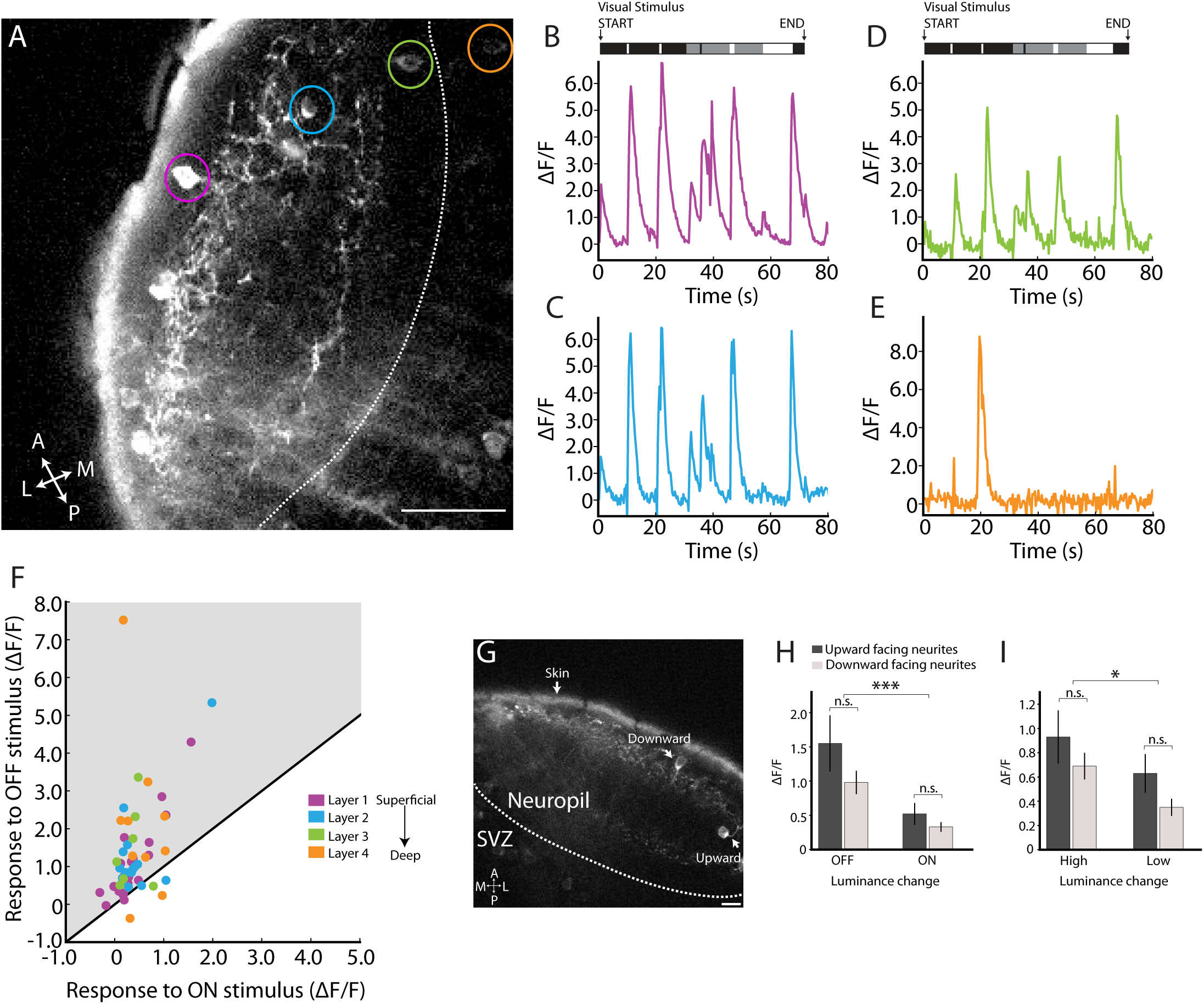
SIN responses to whole-field luminance changes are consistent across neuropil layers. (**A**) Imaging plane in a 7 dpf *Gal4s1156t*, *UAS*:*GCaMP6s* larva. (**B**) Example of a superficially positioned SIN (here denoted as “layer 1”, magenta and indicated by magenta circle in A), demonstrating a characteristic response to all luminance transitions. (**C**) Example trace of a SIN with its cell body positioned in the intermediate layers of the tectal neuropil (here denoted as “layer 2”, blue and indicated by blue circle in A). (**D**) Example trace from a SIN with its cell body positioned in the deepest layer of the neuropil (here “layer 3”, green and indicated by green circle in A). Both intermediate and deep SINs show characteristic responses to the whole-field luminance stimuli. (**E**) PVNs labelled in the *Gal4s1156t*, line show different response profiles to whole-field luminance transitions. An example PVN (“layer 4”, orange and indicated by orange circle in A) trace is shown here. (**F**) Most SINs across all neuropil layers (same cells as shown in Figure 1 plus 10 PVN (“layer 4” cells), n = 57 cells in 17 larvae) show greater responses to stimulus darkening versus brightening. (**G**) SINs with upward facing neurites (towards the superficial tectum) and downwards facing neurites (into the neuropil) are also observed in the *Gal4s1156t* line. (**H, I**) No statistically significant differences in darkening, brightening, high and low luminance stimuli are seen between populations of SINs with upward or downward oriented neurites. However differences between darkening vs. lightening and high vs. low luminance responses are maintained in both populations (unpaired t-tests for upward orientated vs downward oriented neurites, p = 0.13 (darkening), p = 0.22 (brightening), p=0.36 (high luminance change), p= 0.08 (low luminance change; one-way ANOVA with Tukey’s correction for multiple comparisons for across condition comparisons, p = 0.0002 (darkening vs. brightening) and p = 0.036 (high vs. low luminance change), same cells used in Figure 1 and Figure 2F). Scale bar in (A) = 20 µm, (I) = 10 µm. A = anterior, P = posterior, M = medial, L = lateral. *** denotes p < 0.0005; ** p< 0.005, * p < 0.05, n.s. = not statistically significant. Error bars are s.e.m.

The bright loom stimulus consisted of a white expanding disc on a gray background and the dark loom stimulus consisted of a black expanding disc on a gray background. Looming stimuli and whole-field flash stimuli were presented binocularly. For RF mapping and luminance step experiments, stimuli were presented monocularly and calcium responses recorded from the contralateral tectum. Whole-field flash stimuli were presented for 0.5 or 1.0 sec with periods of full black, white or gray screen presentation for 4 s or 10 s following flashes.

### Luminance step stimuli

Luminance steps were generated using custom scripts for PsychoPy (Peirce, 2008). Lux values for luminance steps were: 2.7 lux (black screen), 10.9 lux, 29.8 lux, 67.2 lux, 92.8 lux (white screen). Each luminance step was displayed for 10 s. For analysis a custom written python script was used to select single cell ROIs, their ΔF/F values calculated and baseline changes in the traces corrected using asymmetric least square smoothing (ALS, Eilers and Boelens 2005). For each luminance step the baseline was calculated as a mean of 2-7 frames before stimulus onset and the response as the maximum value in the first 5 frames after the luminance change (baseline subtracted). The stimulus was presented twice and an average value from the two repetitions was calculated.

### Receptive Field Mapping

Receptive Fields were mapped using a checkerboard stimulus with 4 x 3 square grids of ∼20° or 10 x 6 square grids of ∼9° of the larva’s visual field (total area 78° x 60°) generated using custom scripts in PsychoPy (Peirce, 2008). For each stimulation, a single square was chosen randomly to flash to white for 1 s, while the background was kept black. Each white square presentation was followed by 6 s of a black screen. The stimulus was designed so that each square within the grid flashed from black to white twice. Analysis was performed using a custom written python script, where single cell ROIs were selected and ΔF/F values with corrected baseline calculated. The receptive field was calculated using the maximum response during the presentation of each grid square and averaged across two trials. In order to calculate the size of the receptive field, the calculated array was interpolated by a factor of 100 for a smoothed fit, the array was thresholded to the half-maximum and the contour diameter calculated. 10 SINs were exclusively tested with RF mapping, 6 cells with both RF mapping and luminance stimuli and 8 cells with luminance stimuli only.

### Electrophysiology

Whole-cell patch clamp recordings were performed on 6-10dpf *Gal4s1156t, UAS:Kaede* larvae. Larvae were anesthetized on ice and embedded in 2% low melting point agarose (Invitrogen) in external solution. To allow for electrode access to the tectum, a small incision was made with a sharp electrode and used to remove the skin above both tecta immediately prior to recordings. SINs were visualized using the GFP filter on the Zeiss Axioscope2. Recording solutions were as follows: External solution (in mM): 115 NaCl, 2 KCl, 10 HEPES, 2 CaCl_2_, 10 Glucose, 1.5 MgCl_2_, pH 7.4; Internal solution (in mM): 110 Potassium gluconate (C_6_H_11_KO_7_), 10 KCl, 5 NaCl, 1.5 MgCl_2_, 20 HEPES, 0.5 EGTA, pH 7.3. Patch pipettes were pulled from Borosilicate glass (O.D. 1.5 mm, I.D. 0.86 mm; Sutter Instruments BF 150-86-10). The resistance of patch pipettes ranged from 8–10 M. Data were low-pass filtered (2 kHz, Axopatch 200B, Axon Instruments), digitized (10kHz, Digidata 1440A, Axon Instruments) and analyzed using pClamp software (Molecular Devices). Visual stimuli during recordings were presented using a miniature OLED screen (eMagin).

## Results

### SINs are robustly tuned to changes in whole-field luminance

To investigate the functional properties of SINs, we used the *Gal4* line in which they were first identified, *Gal4s1156t* (Del Bene et al., 2010). This line was discovered in an enhancer-trap screen using a basal promoter linked to the gene encoding the transcriptional activator Gal4-VP16 (Scott and Baier, 2009; Scott et al., 2007). To characterize SIN response properties we measured changes in calcium as a indictor of neural activity by crossing the *Gal4s1156t* driver line to a *UAS:GCaMP6s* sensor line. Consistent with previous reports, we identified strong responses to whole-field flashes in SINs in the *Gal4s1156t* line (Figure 1). Expanding on the stimulus sets employed previously (Del Bene et al., 2010; Dunn et al., 2016; Preuss et al., 2014), we presented flashes with a range of six moderate-to-strong luminance changes across the whole field (i.e. white to gray, gray to black, gray to white, black to gray, white to black, and black to white). To exclude the confounding influence of retinal light adaptation on response properties, we kept the overall light levels constant (ca. 0.2 lux) just prior to and during imaging sessions. At this larval age, zebrafish rely exclusively on cone photoreceptors for vision – rods become functional only several days later (Branchek, 1984; Branchek and Bremiller, 1984). We performed behavioral experiments at the same time every day, in the early afternoon of the larva’s subjective day, to account for the well-documented circadian oscillations in light sensitivity (e. g., Moore and Whitmore, 2014).

We found that any whole-field luminance step tested was sufficient to drive responses in SINs (Figure 1A, B) and this response profile was highly stereotyped across all SINs tested (Figure 1, n= 47 SINs). We tested several flash durations (0.5, 1, 4, and 10s) as shown in Figure 1A, B. Despite consistent responses to all luminance transitions, we observed differences in the amplitude of the responses when the stimulus was darkening (OFF) or brightening (ON). For OFF stimuli (i.e. gray to black, white to gray or white to black), we observed significantly greater responses to the whole-field flash than for ON (i.e. gray to white, black to gray, black to white) (Figure 1C, unpaired t-test, p < 0.0001). In addition to a preference for OFF stimuli, we also observed greater responses when the luminance step was large (i.e. black to white or white to black compared to white and black to and from gray) (Figure 1D, unpaired t-test, p = 0.003). Finally, we tested if baseline activity changed during periods of sustained luminance and found that +F/F values were not significantly different between sustained presentations of a gray, black and white screen (Figure 1E, one-way ANOVA with Tukey’s correction for multiple comparisons, p = 0.468). This suggests that SINs become active in response to transient changes in luminance with an increased response to light decrements.

### SINs receive increased excitatory synaptic input in response to OFF steps in whole-field luminance

To corroborate the calcium imaging data with electrophysiology, we carried out voltage clamp recordings from SINs, visualized in transgenic fish in which SINs were fluorescently labeled for better targeting with the patch electrode (*Gal4s1156t*, *UAS*:*Kaede*). Across 7 cells in 7 individual larvae, we observed repeated membrane depolarizations following an offset of the light stimulation that were sustained for the duration of the lights off condition. The total current from these excitatory postsynaptic currents (EPSCs) was greater for whole-field OFF flashes than for ON flashes, except for one cell where the ON and OFF currents were approximately equal (Figure 1F, G). This result is consistent with the increased calcium responses observed during stimulus darkening in our GCaMP6 imaging experiments.

### Responses to whole-field luminance steps do not vary with position in the neuropil

In the *Gal4s1156t* line, SIN cell bodies are observed in all layers of the neuropil (Figure 2A). Additionally, a subset of neurons within the cell-body layer of the tectum, the stratum periventriculare (SPV), are also labeled in this line. We investigated to what extent, if any, response profiles to whole-field luminance transitions varied across the layers of the neuropil. For this analysis, we assigned each of our 47 SINs (from Figure 1) and 10 *Gal4s1156t*+ periventricular neurons (PVNs), as a control, to one of four tectal subdivisions, roughly corresponding to the stratum opticum (SO) and superficial stratum griseum superficiale (upper SFGS), the deep stratum griseum superficiale (lower SFGS), the stratum griseum centrale (SGC) and the SPV. (We decided not to resolve the layers further without knowledge of additional lamina-specific markers.) We assigned 25 SINs to the upper SFGS (“layer 1”), 15 SINs to the lower SFGS (“layer 2”), and 7 SINs to the SGC (“layer 3”). The 10 PVNs were all located in the SPV (“layer 4”). We observed consistent responses in all three subdivisions of the neuropil (Figure 2A-E). Regardless of layer, most SINs showed greater calcium responses to OFF vs. ON stimuli (Figure 2F). No differences in sustained responses to presentation of whole-field gray, white or black stimuli were observed between the three neuropil layers (data not shown, two-way ANOVA, p = 0.391). The PVN controls, on the other hand, showed more diverse responses (Figure 2E, Sup.Fig.1).

In some cases, we observed SINs with neurites extending upward, i. e., toward the superficial surface of the tectum (Figure 2G). We found no significant differences in the whole-field flash responses between SINs in the neuropil regardless of an upward or downward orientation of their neurites (Figure 2H,I). Tuning of upward-facing SINs was very similar to that of the classical, downward-facing SINs, including preference for OFF and high vs. low luminance changes (Figure 2H, I, for comparisons between upward and downward facing SINs we used unpaired t-tests, p = 0.13 (darkening, OFF), p = 0.22 (brightening, ON), p = 0.36 (high luminance change), p= 0.08 (low luminance change; for comparisons between ON and OFF stimuli and high and low luminance changes regardless of neurite orientation we used a one-way ANOVA with Tukey’s correction for multiple comparisons, p = 0.0002 (darkening vs. brightening) and p = 0.036 (high vs. low luminance change), n= 12 upward-facing SINs, n = 29 downward-facing SINs, from 17 fish, same cells as analyzed in Figure 1 and Figure 2F).

### SINs are specifically tuned to a range of luminance steps, but not to looming

We next asked how sensitively tuned the SINs were to detecting specific luminance changes. To probe SIN luminance sensitivity, we generated a new visual stimulus with incremental luminance changes. Steps of intermediate gray values were presented to *Gal4s1156t* SINs expressing GCaMP6s (Figure 3A). The full set of luminance values extended from 2.7 to 92.8 lux (measured on the surface of the screen). Interestingly we observed SINs with narrowly tuned responses to small luminance steps, intermediate steps and large luminance steps, encompassing the whole range of luminance increments and decrements tested (Figure 3B,C).

**Figure 3:**
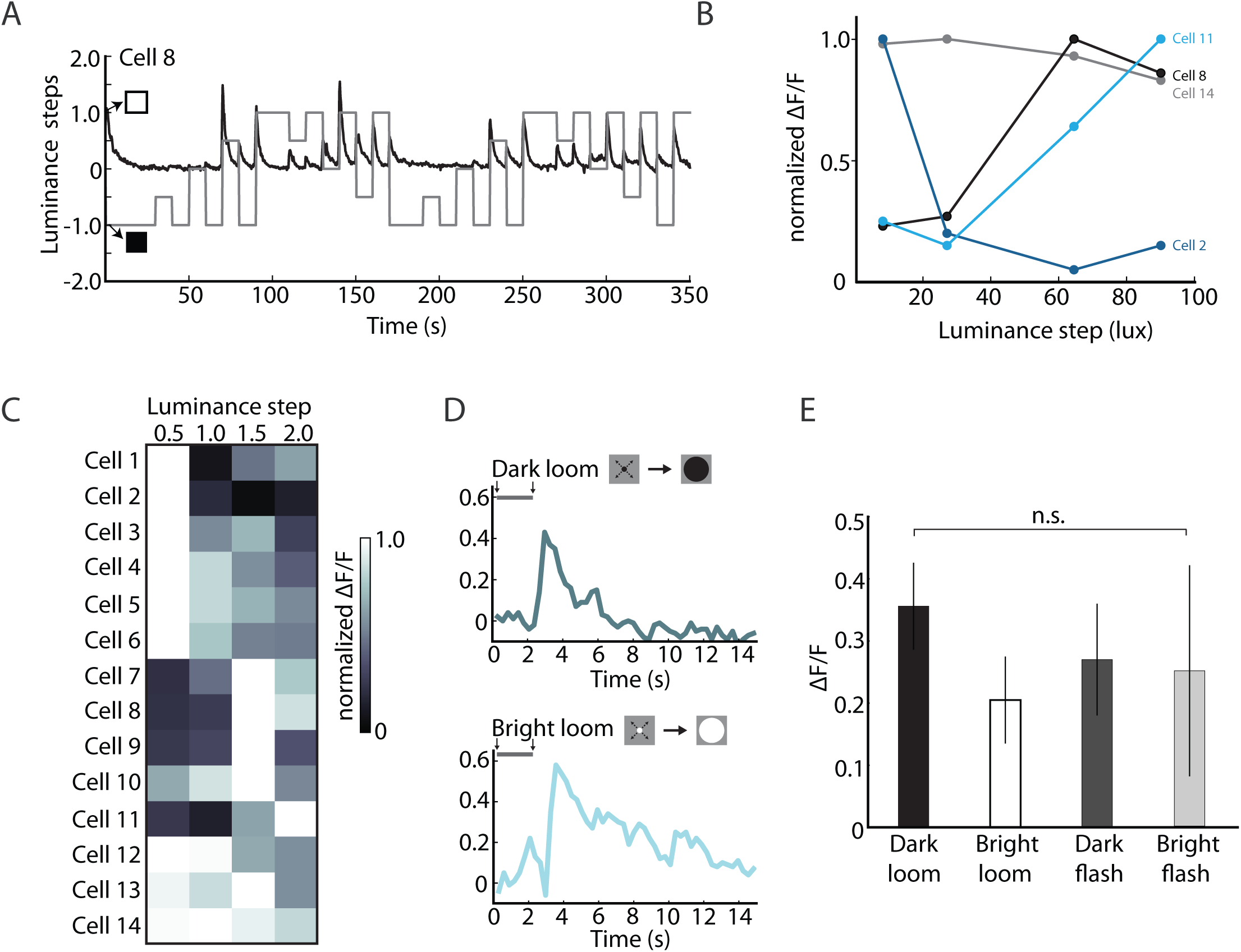
SINs show diverse response profiles to incremental luminance changes. (**A**) SIN responses to finer luminance transitions were tested with a luminance step stimulus. Four progressive steps in whole-field luminance from black (2.7 lux) to white (92.8 lux) were presented in both directions (darkening and brightening). Here responses from an example SIN (Cell 8 in C) in an 8dpf *Gal4s1156t*, *UAS*:*GCaMP6s* larva are shown. Luminance steps are indicated in gray, SIN calcium responses are shown in black. (**B**) Luminance step tuning curves in 4 SINs demonstrate peak tuning distributed across the range of luminance steps tested (n= 14 cells, 8 larvae). The tuning curve for the example SIN shown in (A) is plotted as the black curve. For this cell the strongest calcium responses are observed at larger luminance steps, peaking with a luminance step of 64.5 lux and decreasing slightly at the largest step size tested (90.1 lux). (**C**) Luminance step responses are shown for all 14 SINs recorded in 8 larvae. Heat map shows normalized ΔF/F values for each cell. Luminance steps of 0.5, 1.0, 1.5, and 2.0 correspond to lux changes of 8.2, 27.1, 64.5, and 90.1 lux respectively. Responses to darkening and brightening steps were pooled. (**D**) SINs respond to bright and dark looming stimuli with similar response amplitudes and dynamics. Responses to dark (top) and bright loom (bottom) are shown for the same cell. The visual stimulus is displayed above each trace. (**E**) SINs do not show preferential tuning to bright or dark looming stimuli when compared to equivalent whole-field flashes (n= 11 cells in 5 larvae, one-way ANOVA, Tukey’s correction, p = 0.787). *** denotes p < 0.0005; ** p< 0.005, * p < 0.05, n.s. = not statistically significant. Error bars are s.e.m.

We tested whether SINs were tuned to more specialized stimuli, such as a looming stimulus which produces an overall luminance change but with defined spatiotemporal characteristics (i.e. expansion of a dark spot on a bright background, or vice versa). This was especially pertinent as a previous study integrated SIN responses to looming stimuli into the construction of a behaviorally relevant circuit model (Dunn et al., 2016). To minimize overall luminance changes we presented white, expanding discs on a gray background and black, expanding discs on a gray background (Figure 3D). Expansion velocity was matched to that previously described to elicit optimal behavioral responses to looming (constant radial expansion of 20°/s) (Temizer et al., 2015). Interestingly, we found no differences in responses to bright or dark looming stimuli when compared to full-field flashes (Figure 3D, E, n = 11 cells, 5 larvae, one-way ANOVA, Tukey’s correction for multiple comparisons, p = 0.787). This suggests that *Gal4s1156t* SINs are generally tuned to luminance steps without regard for their spatiotemporal characteristics.

### SIN receptive fields are large

In order to assess the spatial expanse over which SINs are responsive, we mapped their receptive fields (RFs) using a classic mapping strategy (as done previously in zebrafish larvae by Smear et al., 2007). We again used double-transgenic fish carrying *Gal4s1156t* and *UAS:GCaMP6s*. A coarse grid, composed of an array of 12 or 60 squares, each filling about 20° or 9° of the larva’s visual field was presented to one eye of an embedded fish larva positioned on the stage of a two-photon microscope (Figure 4, Sup.Fig. 2). Individual squares were brightened in random sequence, and the evoked calcium responses of SINs were imaged in the contralateral tectum. We found that all SINs had large RFs (23° or greater, Figure 4F). 10 of 16 SINs mapped had contiguous RFs in the range of 40-55° (Figure 4C,D). Five SINs displayed irregular RFs, with foci of peak activity located in spatially incoherent pixels of the grid (Figure 4E). The two cells tested with a 60-square grid (with each square covering ∼9° of the larva’s visual field) showed RFs similar in size and position to those mapped more coarsely (Figure 4A, B). We did not observe any correlation between RF size and the position of the SIN cell body within the neuropil layer (Figure 4G). Together these studies indicate that SINs generally sample over a large portion of the visual field.

**Figure 4:**
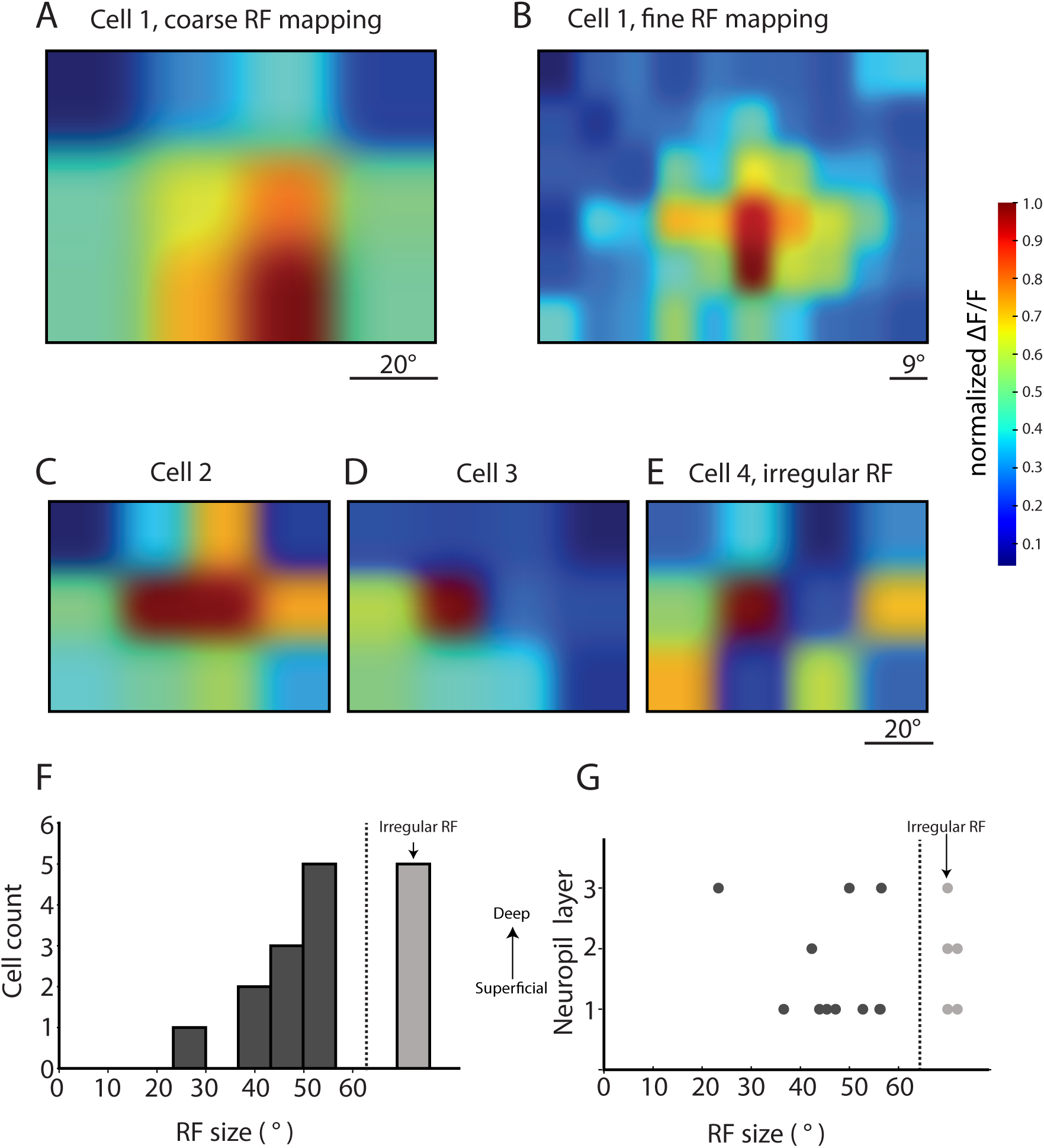
The majority of SINs have large receptive fields. Receptive fields of SINs were mapped using a 12-square grid stimulus. White squares corresponding to 20° of the larva’s visual field were flashed over the whole visual field on a black background. (**A**) An example SIN RF. (**B**) Finer mapping using a 60-square grid for the same cell in (A) revealed a similar RF map. (**C-E**). Three additional examples of SIN RFs mapped with the 12-square grid stimulus. Most cells had a central area of maximal activity (C, D), approximately one-third of cells had an “irregular” RF with areas of highest activity that were not contiguous (E). (**F**) For SINs with contiguous RFs, RF areas were calculated and plotted. (**G**) SINs with different RF sizes were found in all neuropil layers (n= 16 cells, 8 dpf and 9 dpf larvae, 8 larvae).

### SINs are molecularly heterogeneous

In addition to *Gal4s1156t*, several molecular markers for SINs have been reported, including the extracellular matrix protein Reelin and the GABAergic markers Gad67 and GABA (Del Bene et al., 2010). We tested to what extent *Gal4s1156t* SINs overlapped with those expressing Reelin by antibody staining (Figure 5A). Reelin signal was found in 38% of the *Gal4s1156t*-labeled SINs examined (s.e.m. ± 14% in 39 cells from 4 larvae); conversely, many Reelin-positive SINs were *Gal4s1156t*-negative. This indicates that *Gal41156t* and Reelin antibody each label a distinct, partially overlapping subsets of SINs.

**Figure 5:**
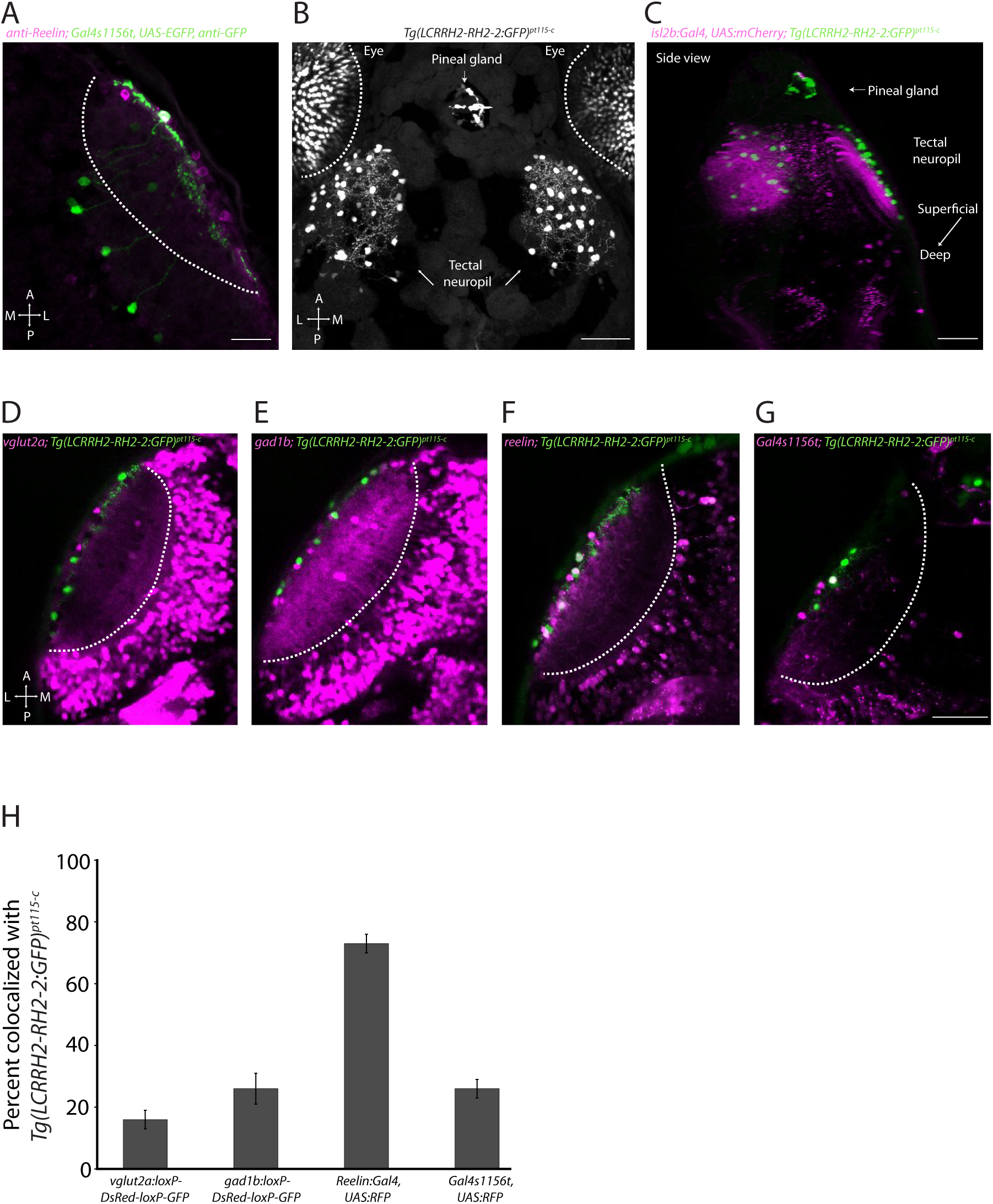
SINs are molecularly heterogeneous. (**A**) Some, but not all *Gal4s1156t*+ SINs are Reelin +. Representative immunostaining of a 8dpf *Gal4s1156t, UAS:EGFP* larva stained with anti-GFP (green) and anti-Reelin (magenta) (n=39 SINs in 4 larvae). (**B**) Top view of a 7 dpf larvae in the *Tg*(*LCR*^*RH2*^*-RH2-2:GFP*)^*pt115-c*^ line. GFP expression is observed in putative SINs as well as in the pineal gland and a subset of photoreceptor cones. (**C**) Rotated confocal volume of a 7 dpf *isl2b:Gal4, UAS:mCherry, Tg*(*LCR*^*RH2*^*-RH2-2:GFP*)^*pt115-c*^ larva shows SIN cell bodies (green) in the *Tg*(*LCR*^*RH2*^*-RH2-2:GFP*)^*pt115-c*^ line are anatomically restricted to superficial layers of the optic tectum (RGC axons labelled by *isl2b:Gal4, UAS:mCherry,* magenta). Putative SINs in the *Tg*(*LCR*^*RH2*^*-RH2-2:GFP*)^*pt115-c*^ line are predominantly localized to the stratum opticum (SO) and the superficial layers of the stratum fibrosum et griseum superficiale (SFGS), consistent with previous SIN morphology (n= 20 larvae, 7 dpf). (**D-G**) Representative examples of GFP+ SINs labelled in the *Tg*(*LCR*^*RH2*^*-RH2-2:GFP*)^*pt115-c*^ line and lines expressing several previously identified SIN molecular markers. (**D**) Crossing the *Tg*(*LCR*^*RH2*^*-RH2-2:GFP*)^*pt115-c*^ ^*line*^ (green) to a *vglut2a*:*loxP-DsRed-loxP-GFP* (magenta) transgenic line, shows a small number of GFP+ SINs express vGlut2a. Some, but not all of the vglut2a + SINs colocalize with GFP+. (**E**) Colocalization between GFP+ SINs and gad1b was assessed by crossing the *Tg*(*LCR*^*RH2*^*-RH2-2:GFP*)^*pt115-c*^ ^*line*^ (green) line to a *gad1b*:*loxP-DsRed-loxP-GFP* transgenic line (magenta). Some but not all gad1b+ SINs showed GFP+ expression. (**F**) Colocalization between GFP+ SINs and the extracellular matrix protein reelin was assessed by crossing the *Tg*(*LCR*^*RH2*^*-RH2-2:GFP*)^*pt115-c*^ *line* (green) line to a *reelin:Gal4, UAS:RFP* transgenic line (magenta). Most GFP+ cells are also reelin+. (**G**) Colocalization between the *Tg*(*LCR*^*RH2*^*-RH2-2:GFP*)^*pt*^115^-^_c_ line and another transgenic line labeling SINs, *Gal4s1156t*, *UAS*:*RFP* was assessed by crossing the two lines. Some but not all GFP+ SINs are also labelled in the *Gal4s1156t* line. (**H**) Percentage of colocalized GFP+ SINs in the *Tg*(*LCR*^*RH2*^*-RH2-2:GFP*)^*pt115-c*^ line with each of the molecular marker lines tested in D-G *(n* = 5 larvae in each condition). Scale bar in A is 20 µm. Scale bars in B, C and G are 50 µm. Larva in A is 8 dpf. Larvae in B-G are 7 dpf. Error bars, s.e.m. A = anterior, P = posterior, M = medial, L = lateral. In A, and D-G the white hatched line indicates the neuropil and stratum periventriculare (SPV) boundary.

We next examined the *Tg*(*LCR*^*RH2*^*-RH2-2:GFP*)^*pt115-c*^ line, in which a regulatory element of the zebrafish *RH2-2* green opsin drives GFP expression (Fang et al., 2013). This opsin promoter line had been previously reported to label only a subset of tectal neurons, along with restricted expression in green cone photoreceptors and the pineal gland (Fang et al., 2013). Intriguingly, the labeled neurons are similar in morphology and position to the SINs (Figure 5B). We crossed the *Tg*(*LCR*^*RH2*^*-RH2-2:GFP*)^*pt115-c*^ line to an *isl2b:Gal4, UAS:mCherry* line, which labels all RGC axons, to highlight the neuropil layers. By crossing these two lines, we confirmed that the majority of GFP+ SINs have cell bodies in the SO and the upper layers of the SFGS (n = 20 fish; Figure 5C). Crossing the *Tg*(*LCR*^*RH2*^*-RH2-2:GFP*)^*pt115-c*^ transgene into the background of lines in which glutamatergic, GABAergic or Reelin-positive neurons are labeled (*vglut2a*:*loxP-DsRed-loxP-GFP*, *gad1b*:*loxP-DsRed-loxP-GFP*, and *reelin:Gal4*) revealed that 16% (± 3% s.e.m.) of these SINs express Vglut2a (Figure 5D,H, n = 5 larvae), 26% (± 5% s.e.m.) Gad1b (Figure 5E, H, n= 5 larvae) and 73% (± 3% s.e.m.) Reelin (Figure 5F, H). In addition, 26% (± 3% s.e.m.) of the SINs in *Tg*(*LCR*^*RH2*^*-RH2-2:GFP*)^*pt115-c*^ were also labeled in *Gal4s1156t* x *UAS*:*RFP* fish (Figure 5G, H). These findings show that SINs are heterogeneous in their gene expression and transmitter choice and that none of the molecular markers available to date offer a complete labeling of the SINs.

### SIN responses to whole-field luminance changes require the retina

Considering that many SINs are labeled in a GFP-transgenic line generated using the regulatory region of a photopigment (green opsin) gene, we asked if their light responses might partially arise from a direct photosensitivity of the SINs. To test this hypothesis, we recorded SIN responses in fish in which both eyes had been removed earlier through enucleation. The operated fish larvae were then presented with the whole-field flash stimuli used previously (Figure 1, 2). In only 3 of 12 SINs from enucleated larvae (n = 3 larvae), some calcium responses were observed (Figure 6). These signals probably originated from spontaneous discharges, as they were not consistent across trials and were not synchronized to the visual stimulus. Responses to flashes in control larvae, with eyes present, often occurred within ∼300 ms (1-2 frames of imaging acquisition) of stimulus offset, whereas, in enucleated larvae, responses occurred, if at all, >3 seconds (>10 frames of imaging acquisition) after the stimulus (Figure 6D). Additionally, we observed no differences in responses to ON vs. OFF, high vs. low luminance, or sustained illumination to gray, black or white full screen stimuli (data not shown, one-way ANOVA with Tukey’s correction for multiple comparisons, p =0.377). Together, these results make it unlikely that the *Gal4s1156t* subpopulation of SINs employ a direct, retina-independent photodetection mechanism.

**Figure 6:**
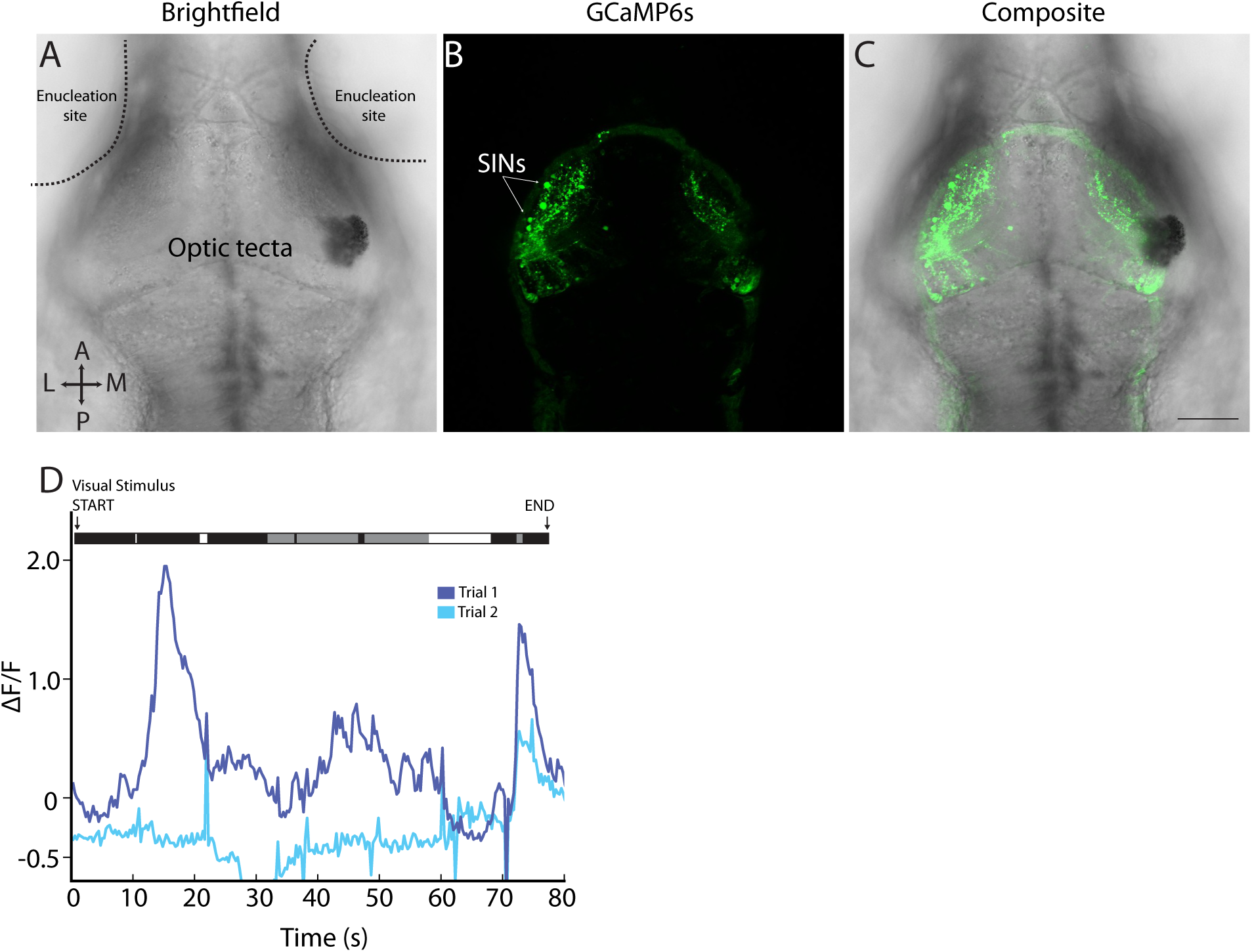
SIN whole-field luminance responses require the retina. (**A-C**) 7 dpf *Gal4s1156t, UAS:GCaMP6s* larva with bilateral removal of eyes. Enucleation site and optic tecta are denoted in the bright field image (A). Following surgical eye removal, GCaMP6s + SIN cell bodies are clearly visible (B). (**D**) Example SIN from the larva shown in A-C. Responses to luminance transitions are not consistent across trials and are not closely synced to the visual stimuli (shown above trace). Scale bar (A-C) = 100 µm.

## Discussion

Can we reconcile the divergent reports of SIN function? Previous studies employed a diverse array of visual stimuli and techniques for labeling and manipulating SINs, making direct comparisons difficult (Barker and Baier, 2015; Del Bene et al., 2010; Dunn et al., 2016; Preuss et al., 2014). While we could not re-test all previously published parameters, we provide the most comprehensive characterization of the *Gal4s1156t* SIN subset to date. We find that these SINs show robust, transient responses to both increments and decrements in light levels, with greatest amplitude responses to OFF stimuli. While, SIN responses to whole-field flashes have been reported (Del Bene et al., 2010; Preuss et al., 2014), we expand this stimulus set by testing incremental luminance changes and show that individual SINs can be tuned to certain luminance step sizes across the entire range of light levels tested. Notably, we also report the first mapping of SIN RFs, which are large and in some cases irregular.

In the SIN population tested, responses to whole-field luminance flashes are largely invariant across all layers of the neuropil and regardless of the orientation of SIN neurites. Layer-specific tuning of retinal inputs for visual stimuli, including size, orientation and direction, have been extensively demonstrated in the larval zebrafish (Dunn et al., 2016; Gabriel et al., 2012; Nikolaou et al., 2012; Semmelhack et al., 2014; Temizer et al., 2015). Moreover, a previous study identified subtypes of SINs with distinct size tuning based on their neurite stratification pattern: superficially stratified SINs preferentially responded to small moving objects, and the more deeply stratified SINs preferentially responded to large moving objects (Preuss et al., 2014). While we have not directly tested the size tuning of *Gal4s1156t* SINs here, our survey suggests that neuropil positional differences do not hold for responses to whole-field luminance changes.

An important caveat with our functional analysis is that it was restricted to the SIN population labeled in the *Gal4s1156t* enhancer-trap line. Attempts to identify a more comprehensive molecular marker for SINs failed. The *Tg*(*LCR*^*RH2*^*-RH2-2:GFP*)^*pt115-c*^ line (Fang et al., 2013) labels many, but not all, SINs. Our study also revealed an unexpected heterogeneity in SIN transmitter use. A previous study demonstrated that *Gal4s1156t* SINs were GABAergic (Del Bene et al., 2010). We confirm that many SINs express the GABA-synthesizing enzyme Gad1b, but report for the first time the presence of Vglut2a+ SINs, an indicator of glutamatergic identity. The lack of complete overlap between molecularly-defined populations of SINs may be due in part to variegation within the *Gal4* lines (i.e. *reelin:Gal4* and *Gal4s1156t*) or the incomplete labeling of glutamatergic and GABAergic populations within the *vglut2a*:*loxP-DsRed-loxP-GFP* and *gad1b*:*loxP-DsRed-loxP-GFP* lines. However, incomplete labeling cannot fully account for the molecular diversity of SINs observed. Rather, we conclude that SINs are more heterogeneous than previously anticipated. A comparison to interneurons in cortex or retina may be instructive, where diverse interneuron populations have been shown to operate in specialized microcircuits (Hangya et al., 2014; Jadzinsky and Baccus, 2013; Masland, 2012).

Extraretinal opsins are present in the zebrafish brain (Fischer et al., 2013), and deep brain photoreceptors have been shown to influence light-mediated behavior (Fernandes et al., 2012). We tested for intrinsic photosensitivity in the SINs by removing the eyes prior to presenting whole-field luminance flashes and recording calcium responses in the SINs. In the absence of retinal input, we observed only a few, apparently spontaneous discharges, which were not synchronized to the presentation of the visual stimulus, making it unlikely that intrinsic photosensitivity contributes to the observed tuning profiles of *Gal4s1156t* SINs.

From where then do SINs receive their tuning for luminance change detection? We propose that a direct retinal synaptic input is likely, although this remains hypothetical. Electrophysiological recordings can give us a clue – we see delays between visual stimulus presentation and onset of activity on the order of 100 ms. In other species, monosynaptic delays in the millisecond range have been reported between retina and tectum, and retina and thalamus (Matsumoto and Bando, 1980; Usrey et al., 1998). However, delays between visual stimulus onset and OFF RGC spiking can be 100 ms in mouse retina (Gollisch and Meister, 2008), similar to the response latencies observed in SINs. One possibility for testing a direct RGC-SIN synapse would require electrical stimulation of RGC axons combined with electrophysiological recordings of SINs, but would likely prove difficult due to the small size of the optic tract and the short distance to the tectum in the zebrafish larva. Better still, electron microscopy reconstruction of the larval zebrafish, which has been performed for other circuits (Helmstaedter et al., 2013; Wanner et al., 2016), should provide a definitive answer regarding the synaptic connectivity of the SINs.

In almost all SINs tested, RFs were substantially larger than 20°, which exceeds reported values for RGC RFs in adult and larval zebrafish (Sajovic and Levinthal, 1982a, 1982b), but is within the range of RF sizes reported for other tectal cell types in both larva and adult (40° ± 4° average RF size reported for 9 dpf larvae and 25-39° in adult zebrafish) (Niell and Smith, 2005; Sajovic and Levinthal, 1982a, 1982b). The spatial layout of SIN RFs is frequently irregular, with several subareas showing peak responses separated by valleys of relative insensitivity. The large size of SIN RFs suggest they may sample over a large area of the visual field, possibly integrating synaptic inputs from multiple RGCs that are scattered over the retinal surface. The identity and functional specificity of these proposed RGC inputs will be an interesting area for future investigation.

A recent study reported SIN responses to different velocities of loom expansion (Dunn et al., 2016). While we confirmed responses to looming stimuli, we did not observe any specific tuning to this type of stimulus. *Gal4s1156t* SINs were equally responsive to bright and dark looming as they were to whole-field flashes. Based on their lack of response selectivity, the *Gal4s1156t* SINs seem inadequate to carry out a specific function in a circuit detecting looming objects. Rather, they might be more generally involved in tasks that require the computation of changes in background luminance levels, perhaps subtracting overall fluctuations in ambient light. However, we cannot exclude the possibility that other SIN populations may contribute to the processing of looming stimuli especially at different temporal frequencies which were not tested here. Our data are consonant with a SIN function in gain control that may prime the tectal circuitry to operate under different luminance conditions. Such mechanisms are extensively utilized in the retina. For example, retinal horizontal cells allow for adaptation to overall luminance levels by acting directly on photoreceptors via feedback inhibition and on bipolar cells via feedforward inhibition (Perlman et al., 1995). In the mouse retina, targeted ablation of horizontal cells resulted in reductions of contrast sensitivity across all spatial frequencies tested in an optomotor assay and deficits in overall visual acuity (Sonntag et al., 2012). Similar gain control and/or background subtraction functions may be important for all levels of visual processing, including the tectum. Such a ubiquitous role may help explain the broad effect of lesioning SINs on visual behaviors (Del Bene et al., 2010, Barker and Baier, 2015). Interestingly, homologs of SINs, or tectal horizontal cells, have been described in the tecta of all vertebrates examined (Guthrie, 1990; Langer and Lund, 1974; Luksch and Golz, 2003; Meek and Schellart, 1978), suggesting a highly conserved and important function in visual processing.

## Acknowledgements

The authors thank Duncan Mearns and Marco Dal Maschio for helpful comments and discussions. We thank S. Higashijima (Okazaki Institute for Integrative Bioscience, Japan) for providing the *vglut2a*:*loxP-DsRed-loxP-GFP* and *gad1b*:*loxP-DsRed-loxP-GFP* fish lines and F. Del Bene (Institut Curie, Paris) for providing the *reelin:Gal4, UAS:RFP, cry:EGFP* fish line.

## Author Contributions

AJB and HB designed research; AJB, TOH and AG performed research and analyzed data; AJB and HB wrote the paper.

